# Ca^2+^ activity is required for injury-induced migration of microglia in zebrafish *in vivo*

**DOI:** 10.1101/2021.11.11.468313

**Authors:** Tian Du, Xi Zhou, Robert Duyang Zhang, Xu-Fei Du

## Abstract

**Objectives:** Microglia are the resident immune cells in the brain. Brain injury can activate the microglia and induce its directional migration towards injury sites for exerting immune functions. While extracellular ATP released from the injury site mediates the directionality of activated microglia’s migration, what endows activated microglia with migration capability remains largely unexplored.

**Methods:** In the present study, we used the larval zebrafish as an *in vivo* model to visualize the dynamics of both morphology and Ca^2+^ activity of microglia during its migration evoked by local brain injury.

**Results:** We found that, in response to local injury, activated microglia exhibited an immediate Ca^2+^ transient and later elevated Ca^2+^ bursts frequency during its migration towards the local injury site (*P* < 0.01). Furthermore, suppression of Ca^2+^ activities significantly retarded microglial migration (*P* < 0.05).

**Conclusion:** Thus, our study suggests that intracellular Ca^2+^ activity is required for activated microglia’s migration.

## Introduction

Microglia are primary immune cells in the brain and derived from a myeloid lineage in the mesoderm [1-3]. Its high sensitivity to ATP and other injury-relevant molecules enables it to rapidly detect and be activated by brain injury and inflammation [4-6]. Besides its activated state for immune functions, intensive studies illustrate microglia at resting state as a multifunction player for regulating diverse neural processes, including synapse pruning [7], myelin formation [8], neuronal activity homeostasis [9], learning [10], forgetting [11], and neuronal death [12,13].

Brain injury can alter microglia status from a physiological resting state to an activated state followed by directional migration towards the injury site [2]. It has been discovered that this process is dependent on injury site-released ATP which is sensed by microglia through P2 receptors, and the concentration gradient of ATP can position the injury site to direct microglia migration [4-6]. However, it is still unexplored what enables activated microglia to migrate. Considering that Ca^2+^ activity is required for the migration of many types of cells [14-18] and Ca^2+^ signaling is important for activating microglia [19,20], we speculate that Ca^2+^ activity may play a role in the migration of activated microglia.

Taking advantage of the optical transparency and small size of the brain in larval zebrafish [21], we performed *in vivo* time-lapse two-photon imaging and visualized the dynamics of both morphology and Ca^2+^ activity of microglia in response to local brain injury mimicked by two-photon laser-based neuron ablation and micropipette stabbing in zebrafish larvae at 5 - 8 days post-fertilization (dpf). Both types of local brain injury could efficiently induce the directional migration of microglia. Interestingly, we found that, during its migration towards the injury site, microglia displayed robust sustained Ca^2+^ burst activities. We then found that the Ca^2+^ burst activities of activated microglia could be suppressed by locally applying 2-Aminoethoxydiphenyl Borate (2-APB), a general antagonist of the Inositol 1,4,5-trisphosphate receptors (IP3Rs) [22]. Importantly, we found that the speed of microglial migration is reduced simultaneously with the suppression of Ca^2+^ activity, suggesting intracellular Ca^2+^ activity is important for microglia migration induced by brain injury.

## Materials and methods

### Zebrafish husbandry and transgenic lines

Adult zebrafish were maintained in an automatic housing system (ESEN, Beijing, China) at 28°C under a 14-hr light:10-hr dark cycle [9]. Embryos were raised up in an incubator. To inhibit pigment formation for two-photon imaging, 0.003% N-Phenylthiourea was added immediately after the collection of embryos. The Tg(coro1a:GCaMP5) line was made by using the same promoter with the Tg(coro1a:eGFP) line which was previously described [23]. The Tg(Huc:NES-jRGECO1a) line was previously described [24].

### In vivo time-lapse two-photon imaging

*In vivo* time-lapse two-photon imaging [25] with a 5 - 10 s interval was performed on the double transgenic larvae Tg(coro1a:GCaMP5);Tg(Huc:NES-jRGECO1a) at 5 - 8 dpf at room temperature (20 - 22 °C). Zebrafish larvae were first paralyzed by α-bungarotoxin and then immobilized in 1.5% low-melting agarose. Imaging was performed under a 25× water-immersion objective (N.A., 0.8) on an Olympus FVMPE-RS-TWIN microscope equipped with a two-photon laser (Newport/Spectra-Physics, USA) tuned to 1000 nm for excitation of GCaMP5. GCaMP5 signal was used to characterize the dynamics of both the morphology and Ca^2+^activity of microglia.

### Two-photon ablation-based local brain injury

The two-photon ablation-based local brain injury was achieved under the Olympus FVMPE-RS-TWIN with an angular tornado-scanning mode. Two-photon laser (800 nm) was targeted on jRGECO1a-positive cells with a circular region of interest (ROI, 5 µm in diameter) on Tg(coro1a:GCaMP5);Tg(Huc:NES-jRGECO1a) larvae. Successful ablation was accepted when the targeted area exhibited a small autofluorescence sphere (5 - 10 µm in diameter) as described in previous studies [4, 26].

### Micropipette-induced local brain injury

Before the injury, the skin on the surface of the hindbrain of zebrafish larvae was removed. Under FVMPE-RS-TWIN, a glass micropipette (1 - 2 µm in diameter of the tip open) made from borosilicate glass capillaries (BF100-58-10, Sutter Instrument) was inserted from the surgery site and the tip was placed around the pretectum area. After insertion, the micropipette was moved back and forth for around 5 - 10 µm to induce tissue damage as previously described [4]. The success of injury induction was confirmed by significant microglial migration to the injury site.

### Local puffing of 2-APB

A stock solution containing 2-APB at 50 mM was first made with DMSO. It was then diluted to 7.5 mM with an extracellular solution (ES) consisted of (in mM): 134 NaCl, 2.9 KCl, 2.1 CaCl_2_, 1.2 MgCl_2_, 10 HEPES, and 10 glucose (pH=7.8), and then loaded into a glass micropipette (1 - 2 µm in diameter of the tip open). Ten minutes after micropipette-induced injury, 2-APB solution contained in the micropipette was ejected out by pulses of gas pressure (4 psi,10 ms in duration,1 min in interval) until the end of imaging. In the control group, exact the same amount of DMSO was diluted in ES and puffed out with the same pulse pattern.

### Data analysis

Analysis of both the Ca^2+^ activity and morphology of microglia was performed with ImageJ (NIH). The image sequences were projected using Z-projection via stacking all slices to the same X-Y plane by maximum grey values. An ROI was drawn to include all the processes and cell body of each microglia. In terms of Ca^2+^ activity, the mean gray-scale value was represented by F, the averaged gray-scale value of the first 100 values from ascendingly sorted F was calculated as F0. The level of Ca^2+^ activity was quantified by (F-F0)/F0 and an increase of > 3 SD was counted as a Ca^2+^ transient activity. The distance to the injury site was measured by calculating the distance between the far side of the microglial cell body and the micropipette tip.

### Statistics

All significance calculation was carried out by Student’s *t*-test. Unpaired or paired Student’s *t*-test was performed for calculating the significance between the data obtained from two groups (migrating microglia *versus* non-migrating microglia, or ES *versus* 2-APB) or from the same group before and after manipulations (before injury *versus* after injury, before ES *versus* after ES, before 2-APB *versus* after 2-APB), respectively. *P* < 0.05 was considered as statistically significant. Data were represented as mean ± SEM.

## Results

### Microglia simultaneously displayed directional migration and Ca^2+^ activities in response to local brain injury in zebrafish *in vivo*

Firstly, we confirmed the effectiveness of the previously described laser-induced and micropipette-induced brain injury for activating microglia. To visualize the dynamics of microglial morphology, we performed *in vivo* time-lapsed two-photon imaging of the double transgenic Tg(coro1a:GCaMP5):Tg(Huc:NES-jRGECO1a) larvae at 5 - 8 dpf at 0.1 - 0.2 Hz (Fig. 1a), in which GCaMP5 (green) and jRGECO (red) are expressed specifically in microglia and neurons of the brain, respectively. Here, GCaMP5 was used to monitor both the morphology and Ca^2+^ activity of microglia, and jRGECO was used for absorbing two-photon laser-associated heat to induce local brain injury in the midbrain where microglia enrich. In response to the local injury, microglia nearby in the same hemisphere became activated as indicated by an amoeboid-like shape and migrated towards the injury site (Fig. 1b). With time, more and more microglia were recruited to surround the injury site (right, Fig. 1b). Furthermore, we used a micropipette to induce local brain injury and observed a similar pattern of activated microglia’s migration but within a smaller spatial range of effect due to more local injury induced by micropipette than two-photon laser (Fig. 1c). This phenomenon was observed in most of the cases examined (20 out of 20 cases for two-photon laser and 13 out of 15 cases for micropipette).

**Fig. 1.**
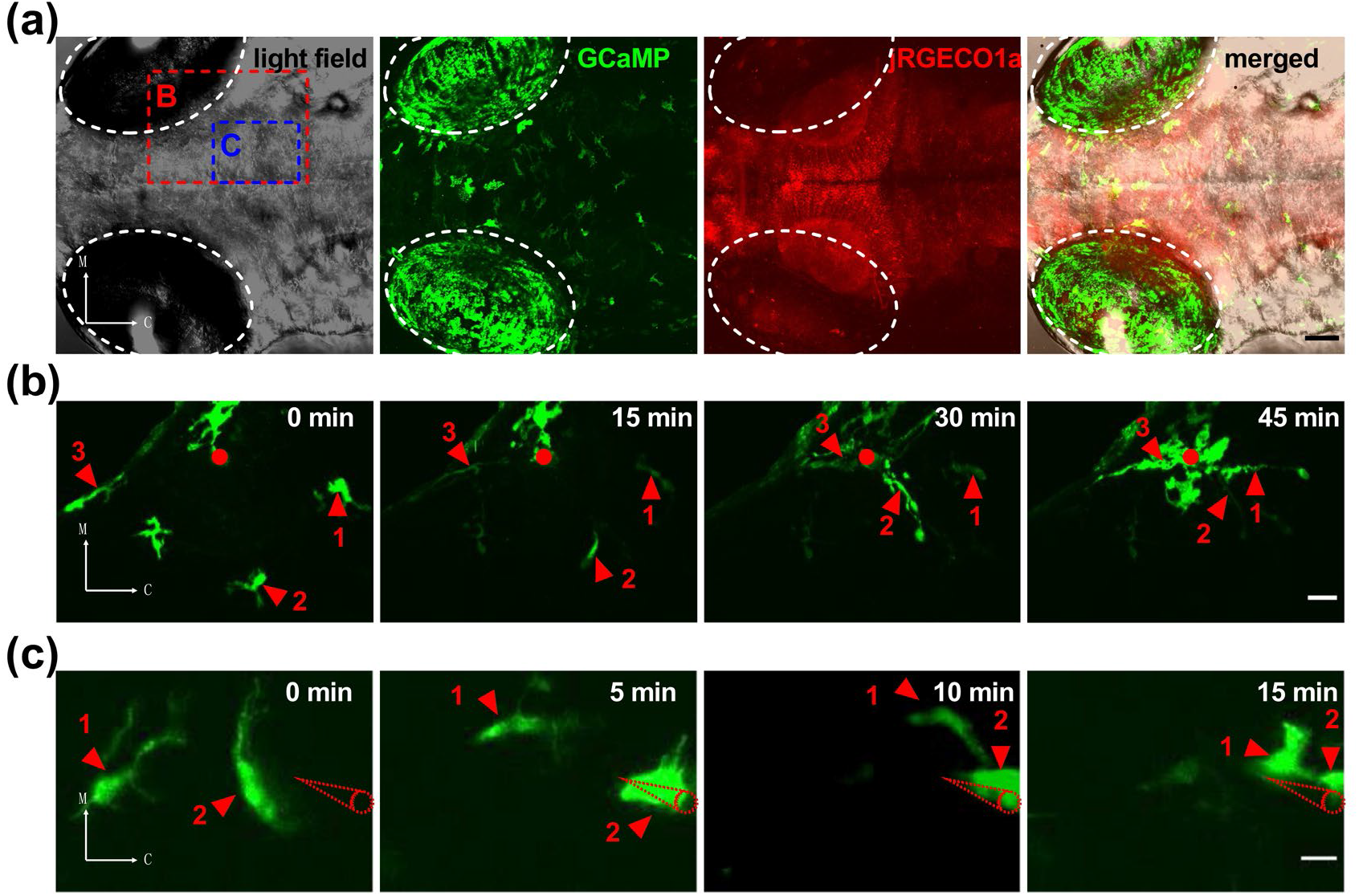
Local brain injury-induced migration of microglia in larval zebrafish *in vivo*1. (a) Whole-brain projection images showing the distribution of microglia (GCaMP-positive, green) and neurons (jRGECO-positive, red) in a double transgenic Tg(coro1a:GCaMP5):Tg(Huc: NES-jRGECO1a) larva at 5 dpf. The green signals on the eyeballs were auto-fluorescence. C, caudal; M, medial. Scale bar: 50 μm. (b) *In vivo* time-lapse (with a 5-s interval) two-photon images showing microglia’s directional migration evoked by two-photon laser-induced local brain injury (red dot). Only time series at a 15-min interval were shown. The time point of 0 min indicates the onset of injury induction. The numbers and arrowheads indicate the microglia traced across the whole imaging. The image field is outlined in (a), and the images were obtained from a 7-dpf larva. Scale bar: 20 μm. (c) *In vivo* time-lapse (with a 4-s interval) two-photon images showing microglia’s directional migration evoked by micropipette (dashed red lines)-induced local brain injury. Only time series at a 5-min interval were shown. The time point of 0 min indicates the onset of micropipette insertion. The numbers and arrowheads indicate the microglia traced across the whole imaging. The image field is outlined in (a), and the images were obtained from a 5-dpf larva. Scale bar: 10 μm.

Then we examined the Ca^2+^ activity of microglia by monitoring changes in GCaMP5 signal (Fig. 2a). A strong Ca^2+^ transient activity was found in nearby microglia immediately after local brain injury and lasted for tens of seconds (arrow, Fig. 2b). And follow-up Ca^2+^ burst activities were observed while those microglia were migrating towards the injury site (“migrating microglia”, red in Fig. 2a, b). Such Ca^2+^ activities were rarely found in those migrating microglia before injury. This type of Ca^2+^ activities were also rare in microglia which located in the contralateral hemisphere of the injury site and did not exhibit directional migration after injury (“non-migrating microglia”, green in Fig. 2a, b).

**Fig. 2.**
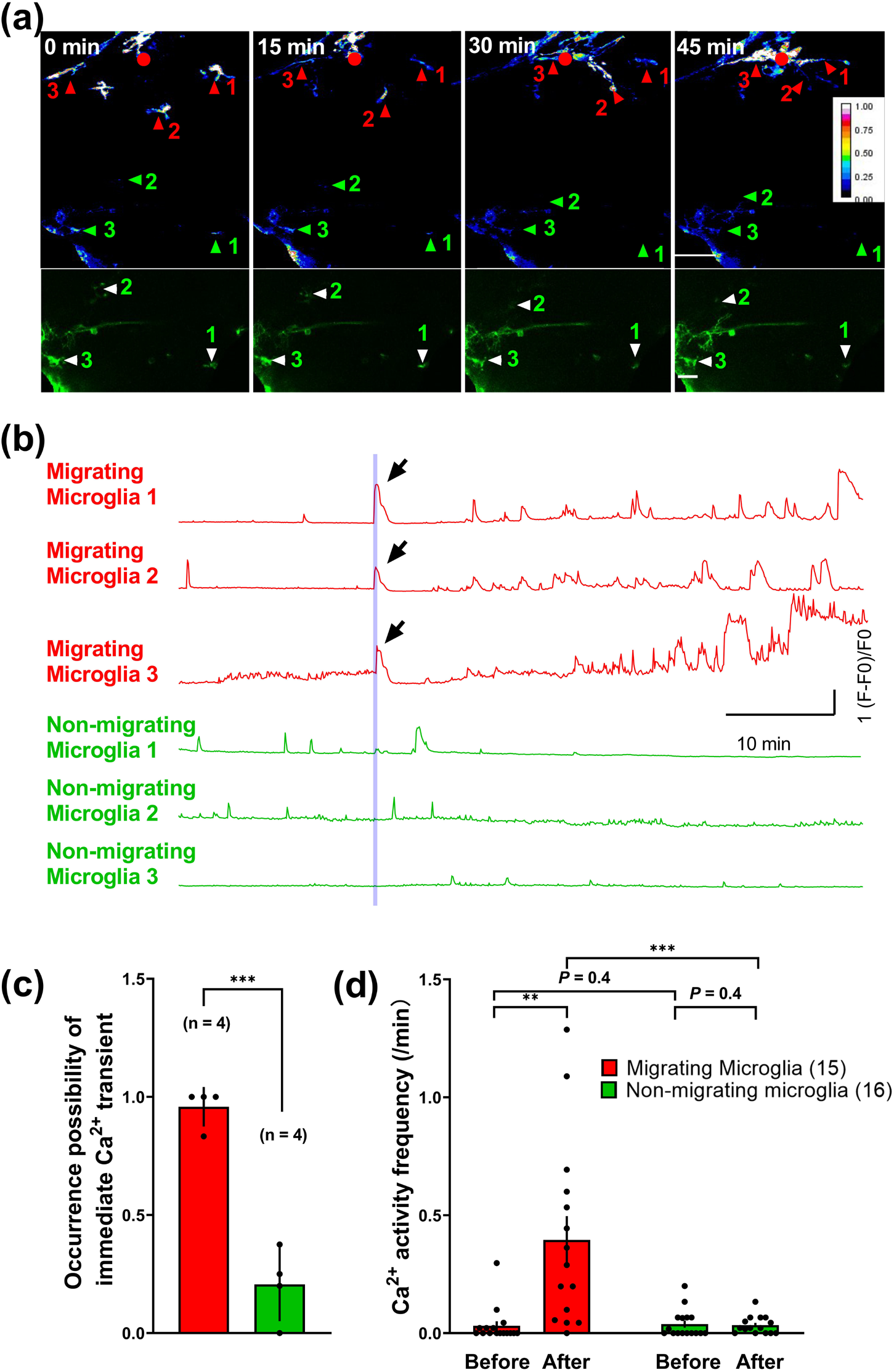
Association between the Ca^2+^ activity and migration of activated microglia. (a) Up: *in vivo* time-lapse two-photon images showing Ca^2+^ activities (in heatmaps) accompanying with microglia migration induced by laser-based local brain injury (red dot) at time 0. The data were obtained from the same larvae with that in Fig. 1B. Red numbers and arrowheads: microglia responsive to the injury; green numbers and arrowheads: microglia located in the contralateral hemisphere and non-responsive to the injury. Down: raw images showing the morphology of non-responsive microglia in the contralateral hemisphere. Scale bar: 50 μm. (b) Traces of Ca^2+^ activities for microglia numbered in (a). The vertical line indicates the onset of local brain injury. Arrow: immediate Ca^2+^ transient induced by the injury. (c) Summary of the occurrence possibility of injury-induced immediate Ca^2+^ transient in migrating (red) and non-migrating (green) microglia. Data were collected from 31 microglia in 4 zebrafish larvae, and each dot on the bar represents the data from an individual larva. (d) Summary of injury-induced later Ca^2+^ activity frequency before and after the local injury in migrating (red) and non-migrating (green) microglia. The number in the brackets represents the number of microglia examined, and each dot on the bar represents the data from an individual larva. Data were collected from 31 microglia in 4 zebrafish larvae. **P < 0.01, ***P < 0.001 (unpaired Student’s *t*-test for (c) and for comparison between migrating and non-migrating microglia in (d), paired Student’s *t*-test for comparison within migrating or non-migrating microglia in (d)). Error bars, SEM

To characterize the association between the Ca^2+^ activity and migration of microglia, we first analyzed the possibility of having immediate Ca^2+^ transients after the injury in both migrating and non-migrating microglia (Fig. 2c). We found that migrating microglia had a significantly higher possibility to have immediate transients in comparison with non-migrating microglia (*P* < 0.001, Fig. 2c). We then compared the frequency of follow-up Ca^2+^ bursts in migrating and non-migrating microglia before and after the local injury induction (Fig. 2d). Both migrating and non-migrating microglia had comparable low Ca^2+^ activities before the local injury (*P* = 0.4). After the injury, the frequency of Ca^2+^ bursts increased markedly in migrating microglia but not in non-migrating microglia (*P* < 0.01 for migrating microglia, *P* = 0.4 for non-migrating microglia, Fig. 2d).

Taken together, these results indicate an association relationship between the migration and Ca^2+^ activity of activated microglia.

### Ca^2+^activity was required for microglia migration towards the injury site

To further explore the role of Ca^2+^ activity in activated microglia’s migration, we examined the effect of Ca^2+^ activity suppression on microglia migration by local application of 2-APB which is a general antagonist of IP3Rs. 2-APB was loaded into the same micropipette used for local injury induction. Just as mentioned in Fig. 2, insertion of the micropipette induced elevated Ca^2+^ activity and directional migration of nearby microglia (“-10 min - 0 min” data in Fig. 3a-d). After 10 minutes of injury-induced migration of microglia, 2-APB was puffed out until the end of imaging. As shown in Fig. 3a, b, after the local application of 2-APB, the level of Ca^2+^ activity in migrating microglia decreased to the baseline. Importantly, the speed of its migration towards the injury site was reduced as well (Fig. 3a, b). In the control, extracellular solution (ES) was ejected out instead of 2-APB with the same pattern. However, after ES application, the microglia still exhibited Ca^2+^ activity and migrated to the injury site (Fig. 3c, d).

**Fig. 3.**
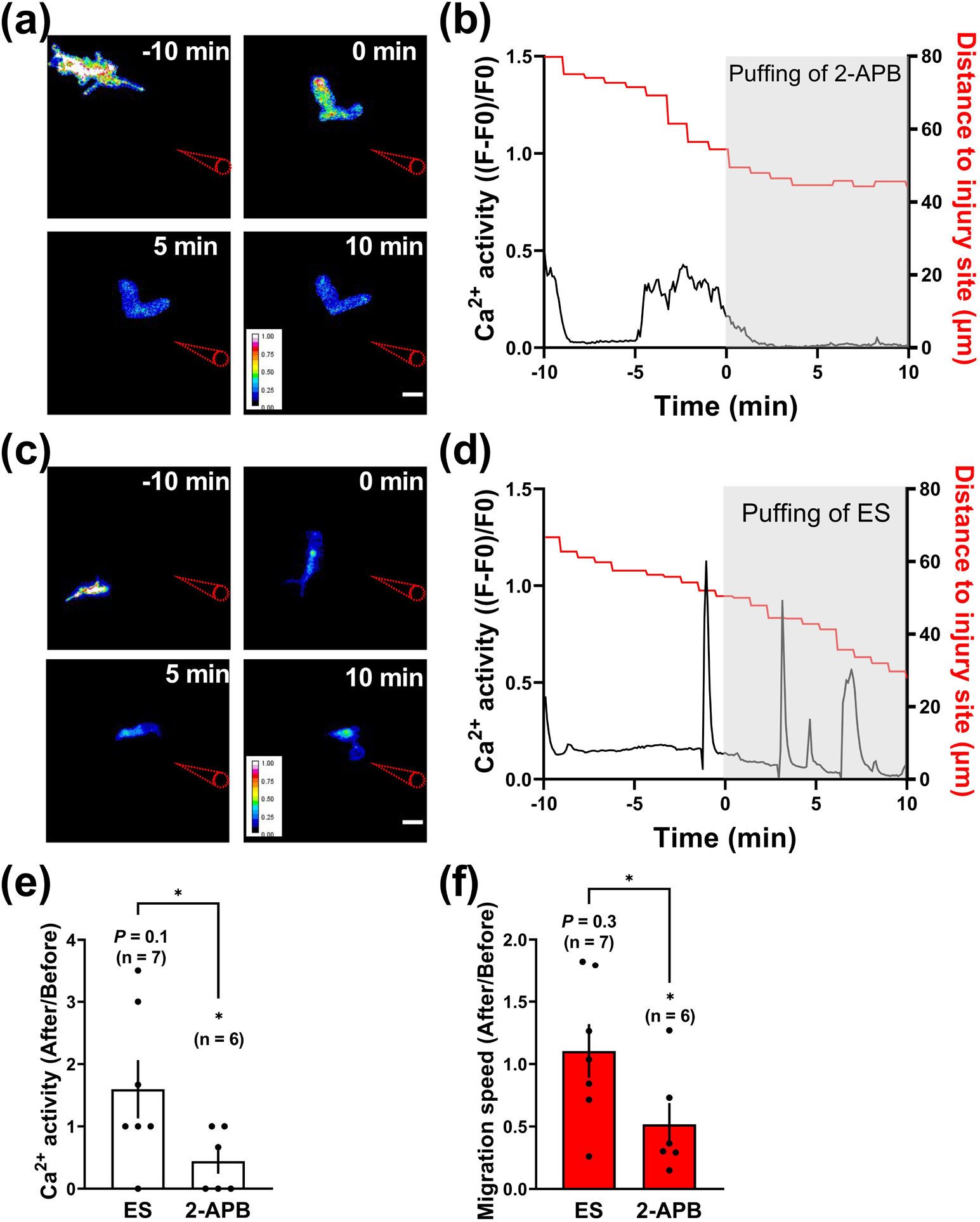
Requirement of Ca^2+^ activity for injury-induced migration of activated microglia. (a) *In vivo* time-lapse two-photon images showing Ca^2+^ activity (in heatmaps) and migration of microglia evoked by micropipette-induced local brain injury (red lines) applied at time -10 min before and after local puffing of 2-APB (just after the time of 0 min). Scale bar: 10 μm. (b) Traces for both the Ca^2+^ activity (black) of the microglia and the distance (red) between the microglia body and micropipette tip for the case in (a). Gray area: application of 2-APB. (c, d) Control case with puffing of extracellular solution (ES) instead of 2-APB. Scale bar in (c): 10 μm. (e, f) Summary of changes in the Ca^2+^ activity (e) and migration speed (f) induced by puffing of ES or 2-APB. Each dot on the bar represents the data from an individual microglia examined. \**P* < 0.05 (paired Student’s *t*-test for comparison within each group before and after the puffing, unpaired Student’s *t*-test for comparison between groups of 2-APB and ES). Error bars, SEM.

To quantify the effects of 2-APB, we calculated and compared the changes of Ca^2+^ activity and migration speed before and after the application of 2-APB or ES. Both the Ca^2+^ activity and migration speed significantly decreased after 2-APB application (*P* < 0.05 for both Ca^2+^ activity and migration speed) but not in the control group (*P* = 0.1 for Ca^2+^ activity and *P* = 0.3 for migration speed) (Fig. 3e, f). These results suggest that intracellular Ca^2+^ activity is required for injury-induced microglia directional migration.

## Discussion

Under the physiological condition, microglia maintain at resting state and locate dispersedly in the brain [1,2]. In response to local brain injury, nearby microglia will be activated and migrate through an ATP concentration gradient produced by injury-induced ATP release [4-6]. Our study reveals an increase of Ca^2+^ activity in migrating microglia as a necessary mechanism for injury-induced directional migration. A previous *in vitro* study reported that isolated mouse microglia displayed spontaneous Ca^2+^ activities [27]. Consistent with our finding, it was reported that Ca^2+^ is highly related to migration in different cell types including endothelial tip cells [14], smooth muscle cells [15], macrophages [16], neural progenitors [17], and oligodendrocyte progenitors [18].

Intracellular Ca^2+^ stores play critical roles in various cell types in regulating Ca^2+^ levels in the cell. Previous *in vitro* studies showed that the intracellular Ca^2+^ store participates in Ca^2+^ activity in microglia and the inhibition of IP3Rs, a widely expressed channel for Ca^2+^ stores, can influence microglia performance in the trans-well assay [28]. As an antagonist of IP3Rs, 2-APB was widely used to inhibit intracellular Ca^2+^ release from the store in various cell types including epithelial cells, platelets, and neutrophils [22]. Our results reveal a significant reduction of injury-induced follow-up Ca^2+^ bursts as well as migration speed after 2-APB application, suggesting the possibility of the involvement of IP3R-associated Ca^2+^ stores in the Ca^2+^ activity of migrating microglia and its important role in mediating microglial migration. As it is reported that 2-APB may also affect multiple TRP channels [29, 30], future investigation is required to confirm the direct involvement of IP3Rs in Ca^2+^ activity of activated microglia *in vivo*. In the present study, we did not exclude the participation of glial cells in the microglial migration, because it is possible that the 2-APB may cause a decrease of intracellular Ca^2+^ concentration in glial cells and result in a reduction of nitric oxide production [31] which could then affect microglial migration [32].

## Conclusion

In summary, our study discovers burst Ca^2+^ activities in activated microglia during its migration to the injury site and suggests that these Ca^2+^ activities are necessary for the directional migration of activated microglia.

## Acknowledgements

We acknowledge Drs. Yu-Fan Wang, Ting-Ting Liu for helpful suggestions, Yu-Xi Li, Sha Li, Yu Qian, Tian-Lun Chen for supporting on imaging experiments, Le Sun for help in image analysis and figure plotting. This research did not receive any specific grant from funding agencies in the public, commercial, or not-for-profit sectors. All authors contribute to this study. TD and XFD designed the study and wrote the manuscript. TD, XZ and RDZ performed the experiments. TD and XZ analyzed the data.

## Notes

**Conflicts of interest:** There are no conflicts of interest.

### Competing Interest Statement

The authors have declared no competing interest.

